# Generating heterokaryotic cells via bacterial cell-cell fusion

**DOI:** 10.1101/2021.09.01.458600

**Authors:** Shraddha Shitut, Meng-Jie Shen, Bart Claushuis, Rico J. E. Derks, Martin Giera, Daniel Rozen, Dennis Claessen, Alexander Kros

**Affiliations:** Origins Centre, Groningen, the Netherlands; Dept. Supramolecular & Biomaterials chemistry, Leiden Institute of Chemistry, Leiden University, the Netherlands; Institute of Biology, Leiden University, the Netherlands; Center for Proteomics and Metabolomics, Leiden University Medical Center, the Netherlands

## Abstract

Cell-cell fusion is fundamentally important for tissue repair, virus transmission, and genetic recombination, among other functions. Fusion has been mainly studied in eukaryotic cells and lipid vesicles, while cell-cell fusion in bacteria is less well characterized, due to the cell wall acting as a fusion-limiting barrier. Here we use cell wall-deficient bacteria to investigate the dynamics of cell fusion in bacteria that replicate without their cell wall. Stable, replicating cells containing differently labeled chromosomes were successfully obtained from fusion. We find that the rate of cell-cell fusion depends on the fluidity of cell membranes. Furthermore, we show that not only the efficiency but also the specificity of cell-cell fusion can be controlled via a pair of synthetic membrane-associated lipopeptides. Our results provide a molecular handle to understand and control cell-cell fusion to generate heterokaryotic cells, which was an important step in the evolution of protocells and of increasing importance for the design of synthetic cells.

## Introduction

The structural and functional complexity of modern bacterial cells evolved gradually over hundreds of millions of years from much simpler enclosed protocells (Szostak, Bartel, and Luisi 2001). These early cells are thought to have resembled self-organizing lipid spheres containing stable catalytic actitivity or primitive metabolism (Monnard and Deamer 2002), but lacking a rigid cell wall. Lipid vesicles are widely used to study the behavior of protocells because they are capable of compartmentalization as well as growth and proliferation (Szostak, Bartel, and Luisi 2001; Adamala and Luisi 2011). Proliferation of such vesicles involves dramatic shape perturbations, such as fission, tubulation, and vesiclulation, which likely preceded the coordinated cell division of modern walled bacteria (Svetina 2009; Hanczyc and Szostak 2004). However, because lipid vesicles are inherently limited in terms of their internal cytoplasmic complexity, consisting of only minimal catalytic components, new models are needed that more closely resemble protocells to effectively study their early evolution (Briers *et al*. 2012; Errington *et al*. 2016). This is particularly needed to examine mechanisms and genetic consequences of cell fusion, an early mechanism of microbial horizontal gene transfer (Kotnik 2013; Soucy, Huang, and Gogarten 2015; Naor and Gophna 2013).

Cell fusion has been studied in many different eukaryotic cell types (Chen *et al*. 2007) and is crucial for tissue repair and regeneration, phenotypic diversity, viral transmission and recombination (Ogle, Cascalho, and Platt 2005). The process of fusion proceeds via several steps: cell adhesion, recognition of cell surface components, membrane remodelling and in some cases nuclear fusion (Zito *et al*. 2016). These processes are highly influenced by lipid-lipid interactions (Chernomordik, Kozlov, and Zimmerberg 1995) which have been studied using coarse grained lipid models and lipid vesicles (Smeijers *et al*. 2006; Marrink and Mark 2003). Fusion in eukaryotic cells is induced via SNARE proteins that form complexes to bridge together membranes by pulling cells close to each other (Hanson, Heuser, and Jahn 1997). The potential for SNARE proteins, or related tools that bridge membranes, to facilitate bacterial fusion have not yet been explored. Studying cell/membrane fusion in eukaryotes and lipid vesicles have unravelled details of the molecular mechanism of membrane fusion; however these systems are highly divergent in terms of cellular and molecular complexity and are not representative of bacterial fusion, which may be common in species lacking a cell wall.

Many bacterial species can transiently shed their cell wall when exposed to environmental stressors like cell wall targeting antibiotics and osmotic stress (Claessen and Errington 2019). When these stressors are removed, wall-deficient cells can rebuild their cell wall and revert to their walled state. Alternatively, prolonged exposure to these stressors can lead to the formation of so-called L-forms, which can efficiently propagate without their wall (Mercier, Kawai, and Errington 2014; Innes and Allan 2001; Glover, Yang, and Zhang 2009; Studer *et al*. 2016). Much like lipid vesicles, L-form growth and division is regulated by physicochemical forces that deform the cell membrane, leading to an irregular assortment of progeny cells. However, L-forms contain the sophisticated machinery of modern cells which is lacking in protocell models based on giant lipid vesicles (Briers *et al*. 2012). This makes them suitable to understanding the dynamics and consequences of cellular fusion, as well as to identify factors that affect this process.

In this study we show that fusion between L-form cells is a dynamic process whose frequency is dependent on the age of the bacterial culture; this, in turn, is determined by the fluidity of the cell membrane, which we confirm by chemically manipulating membrane fluidity. In addition, we demonstrate for the first time that complementary lipidated coiled coil lipopeptides (structurally similar to SNARE proteins) increase the efficiency and specificity of cell-cell fusion. Importantly, fusants resulting from this process are viable and express markers from both parental chromosomes. This opens up avenues to design complex heterokaryotic/hybrid cells that have potential not only to answer questions on evolution of complexity but also enable novel applications in biotechnology.

## Results

### A dual marker system for identifying cell-cell fusion

In order to study cell-cell fusion, we created two fluorescent strains by integrating plasmids pGreen or pRed2 into the *attB* site in the genome of an L-form derivative of the actinobacterium *K. viridifaciens* (Fig. 1A). The strain carrying pGreen constitutively expresses EGFP and is apramycin resistant, while the strain carrying pRed2 constitutively expresses mCherry and is hygromycin resistant (Fig. 1A). We first confirmed resistance to these antibiotics by determining the susceptibility of each strain to both antibiotics (Fig. 1B, supplementary fig. 1A). The strain expressing resistance to apramycin (referred to as AG [for Apramycin-Green]) was able to grow at 50 μg mL^−1^ apramycin. The strain that was hygromycin resistant (referred to as HR [for Hygromycin-Red]) could grow at 100 μg mL^−1^ hygromycin. Resistance to one antibiotic did not provide cross-resistance to the other. Confirmation of the fluorescence reporters was obtained via microscopy with cytoplasmic eGFP detected in the AG strain and mCherry detected in the HR strain (Fig. 1C).

**Figure 1.**
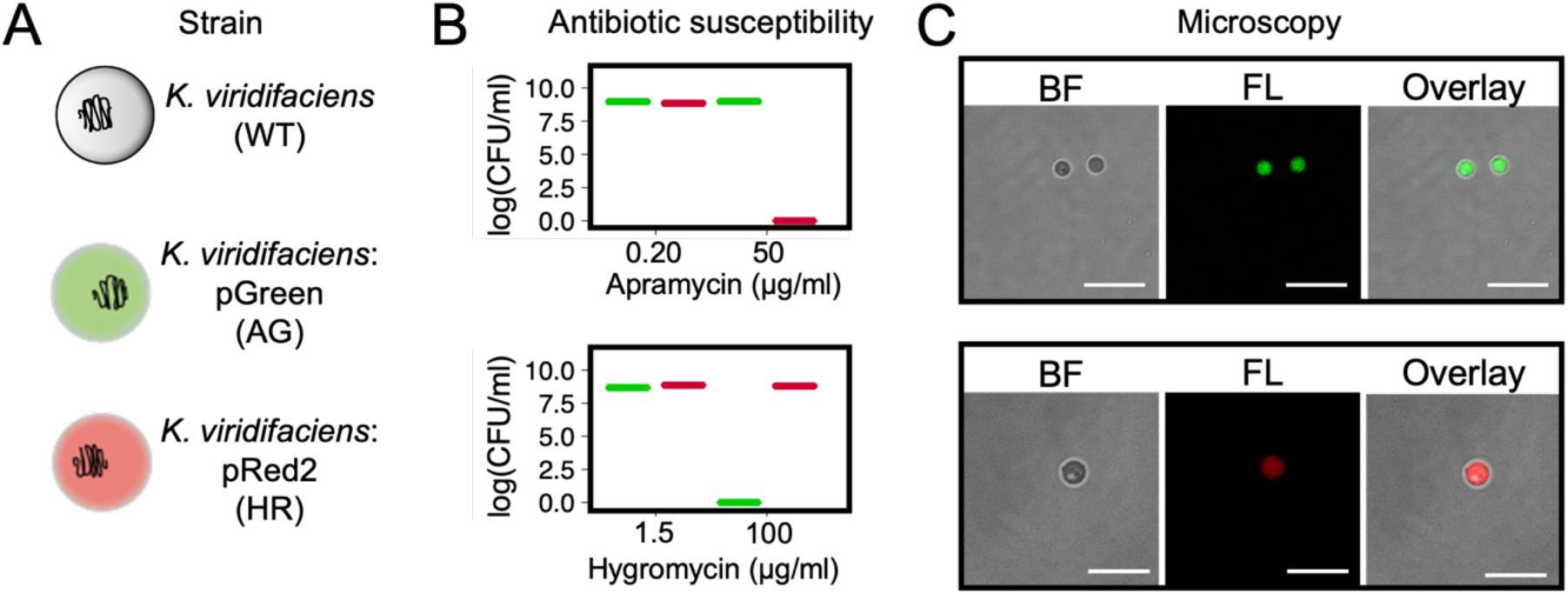
L-forms used in the study. (**A**) The wildtype *Kitasatospora viridifaciens* delta L-form strain was genetically modified to either express apramycin resistance and green fluorescence (AG) or hygromycin resistance and red fluorescence (HR). Each reporter pair (antibiotic resistance+ fluorescence gene) was introduced via a plasmid using the ϕC31 integration system. (**B**) Antibiotic susceptibility testing showed growth of the desired strain at 50 μg/ml apramycin for AG and 100 μg/ml hygromycin for HR. (**C**) Visual confirmation of fluorescence reporters using microscopy indicated a positive signal in the green channel for AG and in the red channel for HR. Scale bar represents 10 μm.

### Fusion of L-form using centrifugation and PEG

L-forms show structural resemblance to protoplasts that are often used for genome reshuffling in plants and bacteria via the process of cell-cell fusion. After fusion these protoplasts can revert back to their walled state. To analyse the ability of L-forms to fuse, we tested some commonly used methods for protoplast fusion (Kieser *et al*. 2000; Baltz and Matsushima 1981; Gokhale, Puntambekar, and Deobagkar 1993) namely, mechanical force induced fusion via centrifugation and PEG-mediated fusion (Fig. 2). Non-specific fusion between AG and HR strains via centrifugation or PEG could result in three different genotypes: AG/HR, AG/AG and HR/HR. However, genetically identical fusants (AG/AG and HR/HR) would not grow on selection plates containing both antibiotics (supplementary fig. 1B). Fusion frequencies determined by growth on both antibiotics are therefore an underestimate of true fusion rates. Centrifuging mixtures of AG and HR at 500 ×*g* resulted in the highest fusion efficiency (1.5 in 10^5^ cells); however, the pellet formed in this case was difficult to handle. Increasing centrifugation to 1000 ×*g* reduced the fusion efficiency to less than 1 fused cell per 10^5^ cells, and no fusion was observed at speeds above 6000 ×g due to cell lysis (Fig. 3A). The fusion efficiency in the presence of PEG was highest at 10 w% PEG with 1 fused cell per 10^5^ cells (Fig. 3B). Higher PEG concentrations, such as 50 w% that is commonly used for protoplast fusion, caused dramatic cell lysis, suggesting that the membrane composition of L-forms is different from protoplasts.

**Figure 2.**
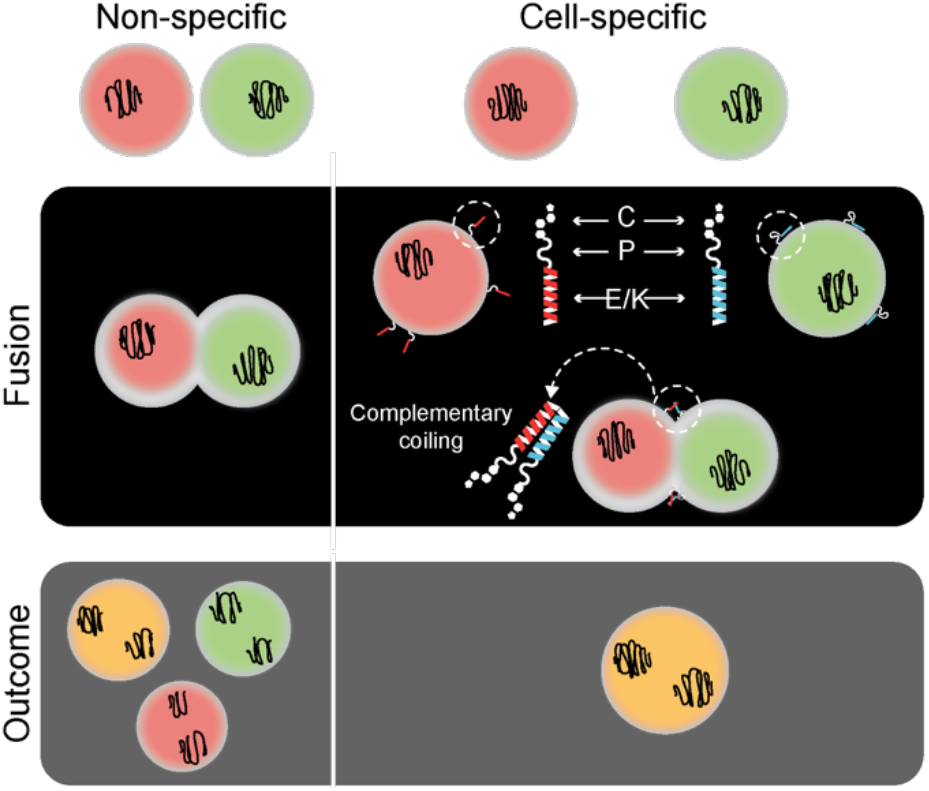
Schematic of L-form fusion. Fusion was obtained by two types of methods: non-specific (centrifugation, poly(ethyleneglycol)-PEG) and cell-specific (coiled coil lipopeptides). The process of fusion (black box) and the outcome (grey box) differs in both cases. For non-specific fusion the membranes come together by dehydration induced by PEG or physical centrifugal force. In the case of coiled coil lipopeptides (CPE and CPK), they dock in the membrane using the cholesterol anchor and pull together opposing membranes upon complementary coiling. This complementarity results in fusion of only oppositely labelled cells unlike that in the non-specific methods.

**Figure 3.**
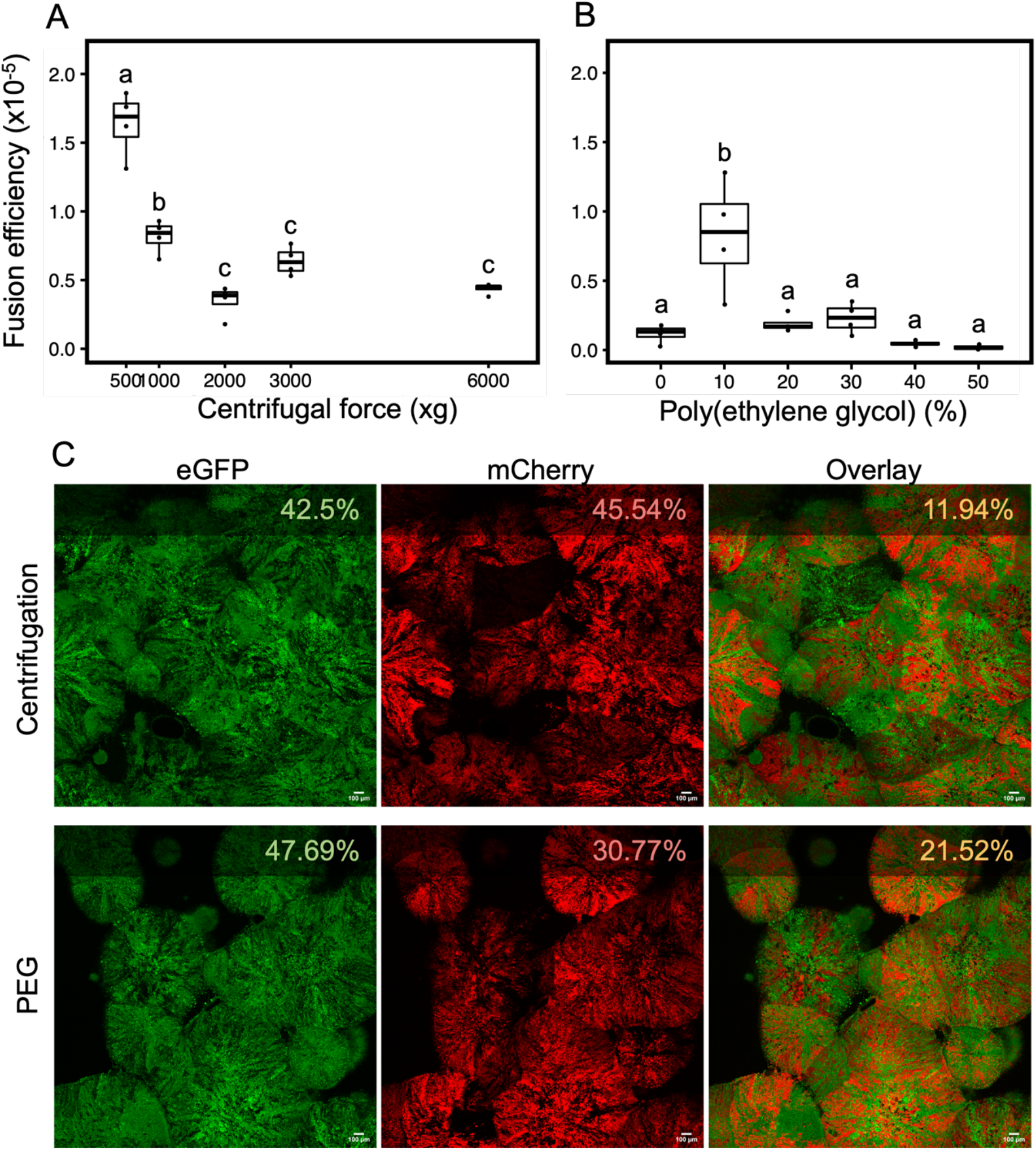
Cell-cell fusion of L-forms. Non specific cell fusion was carried out using either a physical method (centrifugation (**A**)) or chemical method (poly(ethyleneglycol) (**B**)). The fusion efficiency was calculated by dividing the total cell count obtained on double selection media with the cell count of individual parent strain (AG or HR). Increasing centrifugal force leads to a decrease in efficiency (one-way ANOVA, f=15, p=9.77×10^−9^, groupwise comparison Tukey’s HSD). Poly(ethyleneglycol) concentrations also affected fusion efficiency (one-way ANOVA, f=22, p=0.033, groupwise comparison Tukey’s HSD) with 10 %w resulting in the highest efficiency of fusion. (**C**) Fluorescence microscopy of colonies on double antibiotic media after fusion via centrifugation (top panel) and PEG 10 %w (bottom panel). Fluorescence expression (EGFP and mCherry) is indicated as percent in the top right corner of each image and was calculated using ImageJ/Fiji. The overlay image (third column) shows the percent or area occupied by both green and red pixels and is slightly higher for PEG induced fusion. Scale bar = 100 μm.

To verify that the cells growing on plates with both antibiotics (supplementary fig. 1B) were true fusants, we used microscopy. A small patch of biomass growing on media with both antibiotics was imaged using fluorescence microscopy (Fig. 3C). The percent of pixels that were double labelled (i.e. containing both green and red emission) was higher for cells that had undergone fusion via PEG (21.52%) compared to centrifugal force (11.92%). These patches of double labelled cells indicate the presence and subsequent expression of both sets of marker (AG and HR). The presence of green and red patches in the colonies can be attributed to the fact that the polyploid L-forms may consist of an unequal ratio of the two chromosome types. An unequal ratio and expression of markers can lead to a predominantly green (more AG than HR) or red (more HR than AG) colony appearance. Taken together these results show that cell-cell fusion of L-forms is possible and that the resulting colonies contain both chromosomes.

### Fused cells are viable and can proliferate

Successful cell-cell fusion events between different L-form strains combines the cytoplasmic contents and genomes of these cells. To study whether these fused cells (*i.e.* fusant) are viable, timelapse microscopy of individual cells was performed. In viable growing L-forms, membrane extension and blebbing takes place first along with deformation of cell shape (Mercier, Kawai, and Errington 2013; Studer *et al*. 2016). This is followed by daughter cell formation which tend to remain attached to the mother cell. Given the non-binary nature of cell division in wall deficient cells it was difficult to track the exact number of daughter cells originating from one mother cell. Using the wildtype L-forms as a reference for cell growth we looked for the same pattern in fused cells which were viable in the presence of both antibiotics. Colonies from a fusion event were inoculated in double selection liquid media to obtain suspended cultures that could be introduced into a 96 well plate for timelapse imaging in an automated microscope. We applied brightfield and fluorescence imaging every 10 min for over a period of 16 hours (Fig. 4). Importantly, the fused L-forms follow the growth characteristics of wild-type/parental strains as evidenced by blebbing and membrane deformation, as well as smaller daughter cells visibly attached to mother cells (Fig. 4, supplementary movie 1). The fusants also show growth upon subculture into fresh medium containing both selection pressures (supplementary Fig. 2).

**Figure 4.**
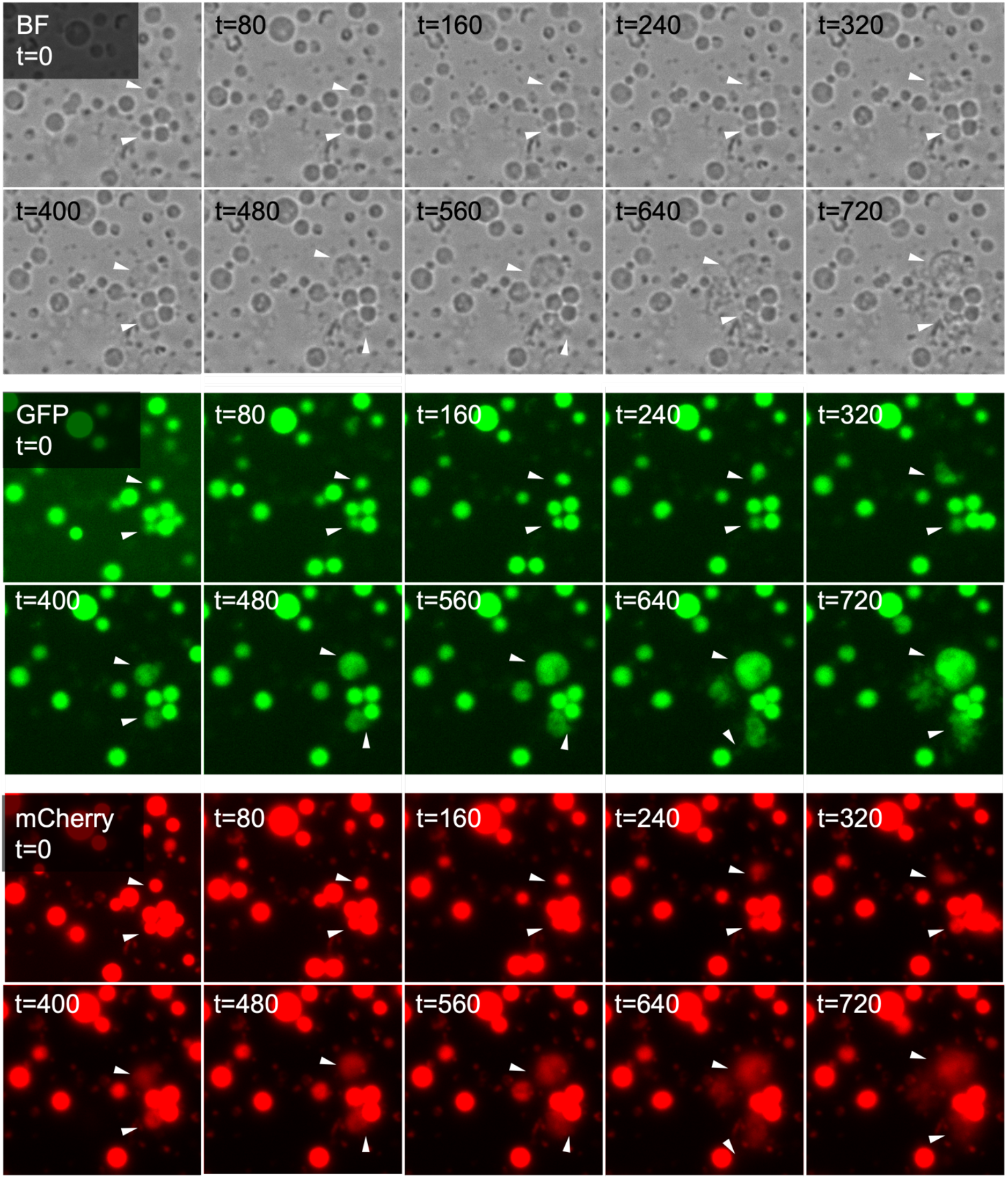
Viability of fused cells. Growth and division of fused cell was tracked over time with brightfield (BF) and fluorescence (GFP and mCherry) microscopy. Images were taken every 10 minutes for a total of 16 hours. The panels (top-BF, middle-GFP, bottom-mCherry) consist of a select few images over this time period (labelled on the top left corner in minutes). White arrows indicate growing cells and membrane extensions. Fused cells also express both fluorescence markers made possible due to cell-cell fusion.

### Membrane fluidity influences fusion efficiency

The bacterial cell membrane largely consists of (phospho)lipids and fatty acids, together with other minor components. The characteristics of these lipids and fatty acids (FA), such as the degree of unsaturation and headgroup composition, determine the physical properties of a membrane. The fluidity of membranes is an important factor governing its fission and fusion ability (Mercier, Domínguez-Cuevas, and Errington 2012; Prives and Shinitzky 1977). Membrane fluidity of L-form cells was quantified as generalized polarization (GP) using the Laurdan dye assay (Scheinpflug, Krylova, and Strahl 2017). This GP value can range from −1 to +1 and inversely correlates to membrane fluidity (*i.e.*, a low GP value indicates a more fluid membrane). Measuring the fluidity for L-forms grown for 1, 3, 5 and 7 days, resulted in a significant GP value increase over time (Fig. 5A, rho=0.732, p=1.87×10^−6^), indicating that membrane rigidity increases as the cultures age. Importantly, this change in fluidity with culture age negatively correlated with the fusion efficiency, as younger cultures fused at twice the efficiency of older cultures (Fig. 5A inset, unpaired t test, p=2.22×10^−6^). To assess the underlying molecular causes for this shift in fluidity, the membrane lipid and FA composition was analyzed using mass spectrometry from L-form cultures of different ages. Over a 7-day period, there was a significant shift in (phospho)lipid/FA composition as the fraction of saturated FAs increased at the expense of unsaturated FAs (Fig. 5B, top panel). This change is consistent with previous reports in *Streptomyces sp*. and *Bacillus sp*. showing that membrane fluidity decreases due to the presence of saturated FAs that stack tightly and thereby make membranes rigid (Mercier, Domínguez-Cuevas, and Errington 2012; Hoischen *et al*. 1997). In addition, the percent of phosphatidylethanolamine (PE) which is known to affect membrane curvature declines with culture age in L-forms. Both factors, an increase in saturated FAs and a decrease in PE, likely underlie the shift in fusion frequency with colony age, although by different mechanisms.

**Figure 5.**
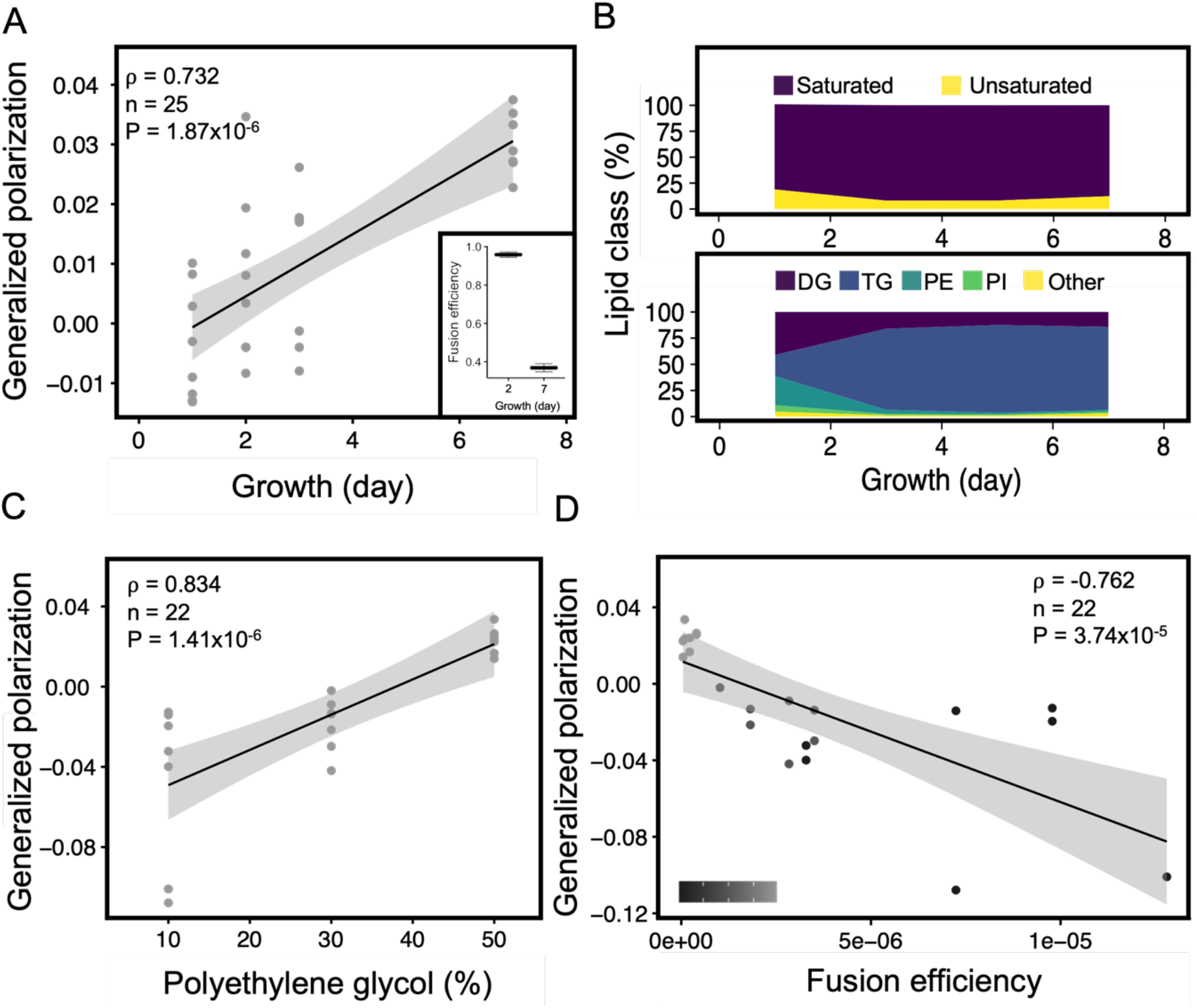
Membrane fluidity affects L-form fusion. (**A**) Fluidity of L-form membranes was quantified as a generalized polarization (GP) value using the Laurdan dye assay. A strong positive correlation was obtained between GP value and the period of growth indicating a decrease in membrane fluidity with increasing culture age (Spearman’s rank correlation test). Age of the culture also has an effect on fusion efficiency (inset, 2 sample t test, p=2.22×10^−6^, n=3) with young 2 day old cultures fusing more efficiently than older 7 day old cultures. (**B**) Analysis of membrane lipids of cultures from different period of growth (1, 3, 5 and 7 day) indicated a change in the percent of saturated and unsaturated fatty acids over time. Specifically the triglyceraldehyde (TG) and phosphatidylethanolamine (PE) show a strong decrease between 1 and 3 day. Both lipids are required for fluidity of the membrane. (**C**) Positive correlation obtained between GP value and the percent of PEG indicating a decrease in membrane fluidity with increasing concentration of PEG (Spearman’s rank correlation test). (**D**) The GP value shows a strong negative correlation with fusion efficiency. A low percent of PEG (10%) leads to slightly more fluid membranes compared to a high PEG percent (50%) resulting in higher fusion (Spearman’s rank correlation test). The grayscale (bottom left corner) indicates PEG percent ranging from 10 to 50.

To causally confirm the impact of membrane fluidity with fusion efficiency, we directly manipulated membrane fluidity by adding PEG into the medium, which is known to induce fusion between two membranes by hydrogen bonding and force adjacent membranes into close proximity via dehydration (MacDonald 1985; Wojcieszyn et al. 1983). When we tested the effect of increasing PEG concentrations on L-form membrane fluidity, we observed a significant positive correlation between GP values and PEG concentrations (rho=0.834, p=1.41×10^−6^) (Fig. 5C) This shows that an increase in PEG leads to reduced membrane fluidity in L-forms. In turn, this caused a decrease in fusion efficiency. Thus a high GP value (i.e. low membrane fluidity) results in low fusion (rho=−0.762, p=3.74×10^−5^) (Fig. 5D).

Taken together these results show that increased membrane fluidity facilitates fusion, which varies naturally during the growth of L-form cells and can be chemically manipulated by the addition of PEG.

### Coiled coil lipopeptides localize to L-form membranes and alter membrane fluidity

PEG-mediated fusion and centrifugation cause non-specific cell fusion and this can result in a low percent of fused cells expressing both EGFP and mCherry (Fig. 2). The recent use of lipidated peptides in cell fusion has shown great promise to improve fusion efficiency, with examples of successful fusion between liposomes or liposomes with various eukaryotic cell lines (Rabe *et al.* 2014; Yang, Shimada, *et al.* 2016; Kong *et al.* 2020; Yang, Bahreman, *et al.* 2016). Coiled coil is a common protein structural motif (supplementary Fig. 3) that contains two or more alpha-helices wrapped around each other to form a left-handed superhelical structure (Koukalová et al. 2018; Robson Marsden and Kros 2010). In previous studies, *de novo* designed coiled coil forming lipopeptides K_4_ and E_4_ were conjugated to cholesterol via a flexible PEG-4 spacer, yielding lipopeptides denoted as CPK_4_ and CPE_4_ (Versluis *et al.* 2013; Zope *et al.* 2013). Using this coiled coil membrane fusion system, efficient liposome-liposome and cell-liposome fusion has been achieved resulting in efficient cytosolic delivery of cargo (Rabe *et al*. 2014; Yang, Shimada, *et al*. 2016; Kong *et al*. 2020). Since L-forms do not posses a cell wall and its outer membrane is structurally similar to (giant) lipid vesicles, we investigated whether coiled-coil lipopeptides CPE_4_/CPK_4_ can be applied to increase the L-form fusion efficiency and introduce cell-specificity. First, we tested whether lipopeptide CPK_4_ could be inserted in the L-form membrane and still form a coiled coil with its binding partner lipopeptide E_4_ (Fig. 2, supplementary fig. 3). Incorporating the CPK_4_ lipopeptide in the membrane allowed docking of the complementary fluorescent labeled peptide E_4_ (fluo-E_4_; Fig. 6A). Docking was also observed when CPE_4_ was incorporated in the L-form membrane, followed by the addition of fluorescent labeled peptide fluo-K_4_. In contrast, no fluorescence was observed when only fluo-K_4_ or fluo-E_4_ was added to L-forms (Fig. 6B). Using image analysis software, we further confirmed membrane localization of the lipopeptide-fluorescent dye conjugate by assessing the fluorescence intensity across the cell along a transect line. A combined plot (supplementary fig. 4) of these intensity values across 10 cells indicates coinciding peaks of fluorescence values of the lipopeptide conjugates with that of gray values of the cell membrane (seen as dark grey rings in brightfield images). The fluorescence intensity on L-form membranes was more distinct when CPE_4_/fluo-K_4_ was used as compared to CPK_4_/fluo-E_4_ (supplementary fig. 4A). Altogether, these results demonstrate for the first time that lipopeptides can be readily incorporated into L-form membranes and serve as a docking point for the complementary (lipo)peptides.

**Figure 6.**
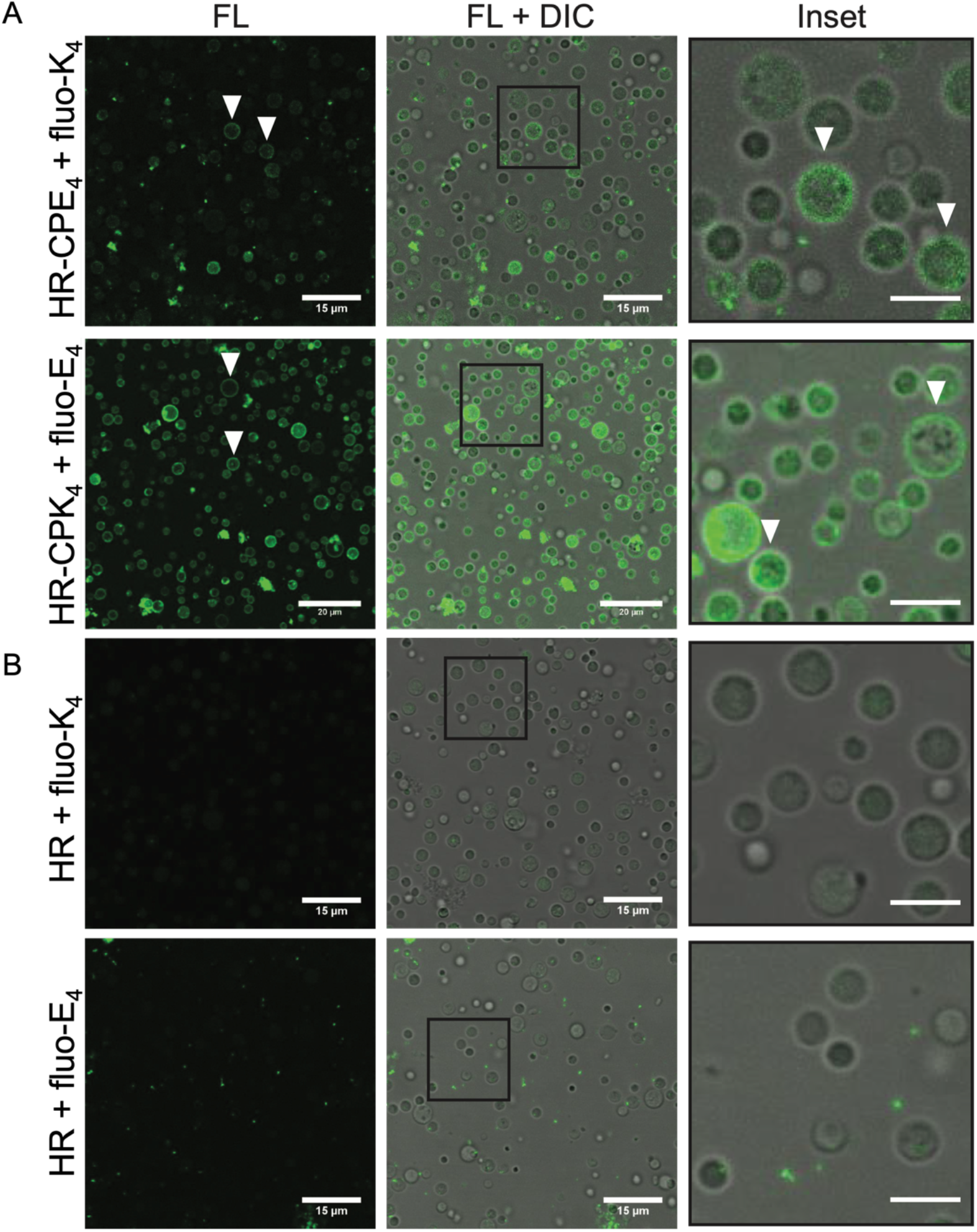
Coiled coil lipopeptides integrate in L-form membranes. (**A**) Confocal microscopy images (fluorescence (FL) and overlay (FL+DIC)) indicating peptide CPE_4_ or CPK_4_ insertion into the L-form membranes and coiled-coil formation with complementary peptides (fluo-K_4_ or fluo-E_4_). White arrows indicate clear membrane insertion. (**B**) In the absence of CPE_4_ or CPK_4_ no binding of the complementary fluorescent peptides (fluo-K_4_ or fluo-E_4_) was observed. Experiments were performed at 30°C, L-forms in P-buffer were incubated with 10 μM of CPE_4_ or CPK_4_ for 30 minutes. Subsequently the unbound peptide was washed via centrifugation and the complementary fluorescent peptides were added. Scale bar = 5 μM.

The incorporation of lipopeptides in L-form membranes prompted us to investigate whether they also influenced membrane fluidity. To test this, L-forms expressing red fluorescent protein (AR and HR strains) were modified with either non-fluorescent labeled CPE_4_ and CPK_4_ so as not to interfere with the emission spectra of the Laurdan dye. The observed GP values reveal that CPK_4_ and CPE_4_ affect the fluidity of L-forms differently. While CPE_4_ decreased fluidity in the AR strain (Fig. 7A), both lipopeptides increased fluidity in the HR strain (Fig. 7B). Interestingly the effect of increased fluidity due to PEG (10 w%) was only observed in the AR strain. These differences in fluidity effects are likely caused by the presence of antibiotics during culturing of the strains prior to the experiment, which are required to avoid contamination in the cultures (supplementary fig. 5). Antibiotics are known to affect membrane fluidity (Bessa, Ferreira, and Gameiro 2018), however the exact mechanism by which they do so is unclear. This inherent difference was observed in the basal GP values of control samples (−0.02 for HR and −0.08 for AR, Fig. 7A-B) as well as in separate measurements for fluidity of strains in the absence and presence of antibiotics (0.01 for HR and −0.10 for AR, supplementary fig. 5). However, all treatments (PEG/lipopeptide) are compared to the control sample of individual strain type; hence, the change in GP value is indeed due to the lipopeptide interaction and the to the presence of antibiotics.

**Figure 7.**
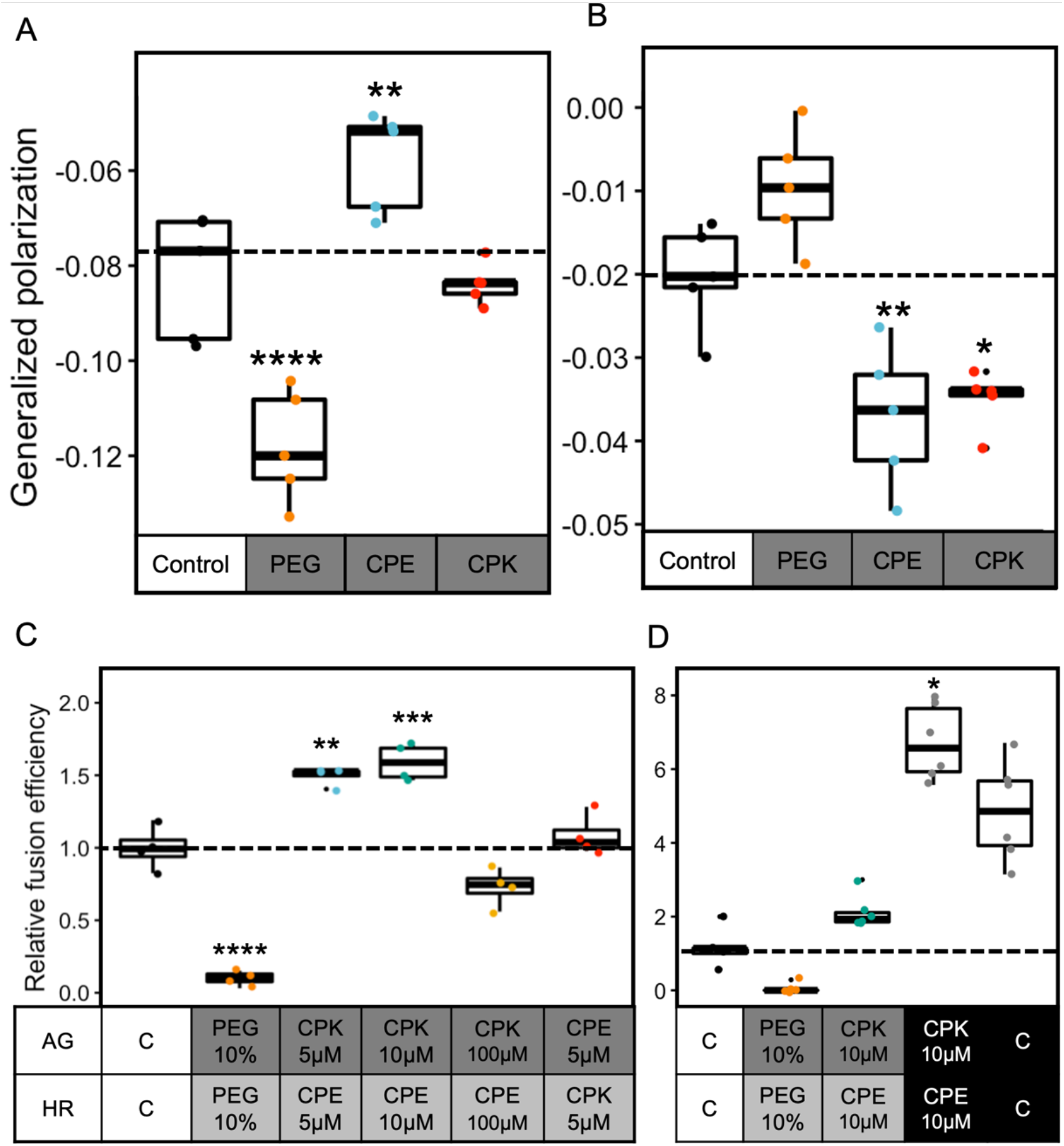
Coiled coil lipopeptides increase membrane fluidity and cell-specific fusion. (**A**) The strain AR shows an increased fluidity on treatment with PEG (p=3.06×10^−6^), a decrease in fluidity on treatment with CPE_4_ (p=2.13×10^−3^) and no change in fluidity with CPK_4_ (One-way ANOVA, F=36, p=4.59×10^−18^ followed by Tukey’s pairwise comparison) compared to the control (dotted line). (**B**) The strain HR shows increased fluidity (low GP value) when treated with CPE_4_ (p=3.11×10^−3^) and CPK_4_ (p=1.4×10^−2^) compared to the control (dotted line) whereas no significant change when treated with 10% PEG (One-way ANOVA, F=36, p=2.83×10^−18^ followed by Tukey’s pairwise comparison). Dotted line is for comparison of GP values to the control where no peptide or PEG was added. (**C**) The AG and HR strains were individually treated with either PEG, CPE_4_ or CPK_4_ at different peptide concentrations to assess the effect on fusion efficiency. Interestingly PEG leads to low fusion despite increasing fluidity because of its non-specific nature. The combination of AG-CPK_4_ and HR-CPE_4_ resulted in highest fusion efficiency relative to the basal level. The increase in relative fusion efficiency is concentration dependent as well as peptide dependent (One-way ANOVA, F=30, p=3.47×10^−14^ followed by Tukey’s pairwise comparison). (**D**) The AG and HR strains were first treated with either PEG, CPE_4_ or CPK_4_. These strains were then directly plated on double selection media in the absence (grey boxes) or presence (black boxes) of 10%w PEG to assess the effect on fusion efficiency. Interestingly PEG leads to low fusion despite increasing fluidity because of its non-specific nature when washed away prior to plating but gives a high efficiency when present during the plating. The treatment with peptides also shows a higher efficiency when in the presence of PEG (Kruskall-Wallis chi-squared = 24.84, p=5.4×10^−5^, followed by Dunnet’s pairwise comparison) compared to the control where no peptide or PEG was added (dotted line).

We next examined how these changes in fluidity affect the process of lipopeptide-mediated fusion. For this, L-form cultures were first adjusted to the same density and split into aliquots. The aliquots were then either untreated (control), treated with PEG or increasing concentrations of the lipopeptide that previously caused an increase in fluidity. HR strains were hence pretreated with CPE_4_ and AG strains were treated with CPK_4_. After treatment for 30 minutes the excess PEG and lipopeptides were removed by centrifugation and the L-forms were resuspended in fresh P-buffer containing DnaseI. The cultures were then thoroughly mixed in a 1:1 ratio, incubated for 30 minutes at 30°C and subsequently plated on selection media for cell quantification. The observed fusion efficiency for each treatment relative to control revealed that treatment of HR with CPE_4_ and AG with CPK_4_ results in a high fusion efficiency as compared to 10 w% PEG or the centrifuged control (Fig. 7C). Furthermore, fusion efficiency was not only dependent on lipopeptide concentration (i.e. decreased fusion at 100 μM) but also on the lipopeptide specificity since AG treated with CPE_4_ resulted in basal level of fusion similar to the control. Higher lipopeptide concentrations also visibly affected cells, causing lysis (data not shown).

Together these results confirm that cell specific fusion of L-forms can be achieved using fusogenic coiled coil lipopeptides.

The two approaches (non-specific via PEG and centrifugation and cell-specific using lipopeptides) used here seem to influence fusion by altering membrane fluidity and bringing membranes together. We then investigated whether combining both fusogens would result in an overall higher fusion efficiency. For this the cells were first treated with the lipopeptides (AG L-forms with CPK_4_ and HR L-forms with CPE_4_) and split into two aliquots. The first aliquot was directly subjected to fusion by mixing the cultures in a 1:1 ratio whereas the second aliquot was mixed and treated with PEG. Here the PEG remained in the environment during the process of fusion. Efficiency calculations showed a 3-fold higher relative fusion in the latter (Fig. 7D) indicating that combining lipopeptides and PEG is optimal for cell-cell fusion. The presence of lipopeptides on the cell surface aids in complementary L-form pairing (AG with HR) bringing the opposing membranes in close proximity, which is an important first step in fusion. Additionaly PEG potentially further reduces the space by membrane dehydration thus facilitating fusion events. Colony imaging further confirmed the presence of more double labelled cells in treatment with PEG (supplementary Fig. 6).

## Discussion

Cell wall deficiency has primarily been studied in the context of stress tolerance and intracellular pathogenicity (Errington *et al*. 2016). The genetic and metabolic modifications required to survive in this wall-deficient state are also being uncovered which has deepened our understanding of their intriguing biology (Glover, Yang, and Zhang 2009; Kawai *et al*. 2019). We here show that wall-deficient L-forms are able to fuse with one another and that membrane fluidity is a key factor influencing fusion efficiency. Additionally, we show for the first time targeted fusion between wall-deficient cells using coiled coil lipopeptides. This opens up avenues for application in the field of biotechnology and the design of synthetic cells.

L-forms are surrounded by a membrane, which are be sufficiently fluid to allow efficient proliferation. *Bacillus subtilis* L-forms that have a defect in formation of branched chain fatty acid (BCFA) suffer from decreased membrane fluidity and as a consequence cannot carry out the membrane scission step (Mercier, Domínguez-Cuevas, and Errington 2012). This phenotype was rescued by supplementing the media with BCFAs in the medium. Less is known about the impact of fluidity on bacterial fusion, although older reports on eukaryotic muscle cell cultures suggest that myoblast fusion was preceded by a decrease in membrane viscosity (Prives and Shinitzky 1977). In this work we showed that the membrane fluidity of *K. viridifaciens* L-forms changes over time. In younger cultures, the fluidity is higher coinciding with the ability of such cells to proliferate efficiently. By contrast, the fluidity decreases in older cultures. The change in fluidity was associated with a change in the ratio of saturated to unsaturated FAs. In our study we found this ratio to be 4.3 for the 1^st^ day of growth which then increased to 11.3 after 3 days (Fig. 5B). Thus the amount of saturated FAs responsible for tighter packing increases over time at the expense of unsaturated FAs. The accumulation of saturated FAs makes the membrane more stiff, which negatively impacts proliferation and fusion efficiency. Notably, compared to protoplasts, L-forms of *Streptomyces hygroscopicus* contained 6 times more anteiso FAs than protoplasts resulting in more fluid membranes (Hoischen *et al*. 1997). Our lipidomics analysis also indicates that L-form membrane composition comprised significant amounts of cardiolipin (CL), phosphatidylinositol (PI) and phosphatidylethanolamine (PE). Both CL and PE are fusogenic headgroups shown to induce fusion between liposomes and extracellular vesicles (Driessen *et al*. 1985), and their the presence may also facilitate L-form fusion.

A pair of complementary fusogenic coiled coil lipopeptides have been previously developed for the targeted delivery of compounds into eukaryotic cells using liposomes. These eukaryotic-liposome models have also been used extensively to understand the process of cell fusion (Daudey *et al*. 2017). For the first time we explored targeted fusion with these synthetic lipopeptides between bacterial cells. Interestingly we observed that the lipopeptides readily insert in membranes of L-forms via a cholesterol anchor (Fig. 6). These lipopeptides remained in the membrane even after several washing steps. The lipopeptide segment of CPK_4_ is known to interact both with its binding partner lipopeptide E_4_ as well as membranes while the lipopeptide E_4_ segment of CPE_4_ does not (Fig. 2). Complementary binding of the lipopeptides brings two opposing membranes in close proximity and ultimately induces fusion (Koukalová *et al*. 2018; Robson Marsden *et al*. 2009). The differences in lipopeptide presentation on the surface can explain the complementarity effect on fusion efficiency of L-forms as well (Fig. 7). Given the ease of lipopeptide docking and subsequent stability on the L-forms, coiled coil lipopeptides provide a promising avenue for studies on targeted compound delivery into wall deficient cells. This may be particularly relevant for L-forms associated with recurring urinary tract infections and potentially mycobacterial infections (Mickiewicz *et al*. 2019; Markova 2017).

The costs and benefits of living as a wall deficient cell depends on the environment. Absence of a protective wall makes them sensitive to changes in osmotic pressure and physical agitation. On the other hand, cells without a wall are resistant to a whole class of cell wall targeting antibiotics (penicillins, cephalosporins), transport to the extracellular space is potentially easier and the cells are stably polyploid. These characteristics can make L-forms a unique model system to study not only cell biology but also questions in the fields of biotechnology, evolution and the origin of life (Briers *et al*. 2012; Errington *et al*. 2016; Shitut *et al*. 2020). The process of cell fusion may have been a mechanism of horizontal gene transfer and species diversification in early life (Küppers and Zimmermann 1983). Understanding this process is hence a key aspect of protocell evolution. L-forms are uniquely suited to replicate these processes thereby providing a mechanistic understanding of the causes and consequences of such fusion. First, the use of coiled coil directed fusion can be extended to synthetic cells to obtain fusions that increase cellular complexity. Second, fusion leads to multiple chromosomes in the same cellular compartment which in turn can result in genetic recombination. Such recombination events can then be leveraged to identify new microbial products and obtain genomically diverse populations of cells. Finally, cell-cell fusion can also help to understand major transitions on the road to increased organismal complexity like multicellularity and endosymbiosis.

## Materials and methods

### Media and growth conditions

All L-form strains were cultured in liquid L phase broth (LPB) and solid L phase media agar (LPMA). LPB consists of a 1:1 mixture of yeast extract malt extract (YEME) and tryptic soy broth supplemented with 10% sucrose (TSBS) and 25 mM MgCl_2_. LPMA consists of LPB supplemented with 1.5% agar, 5% horse serum and 25 mM MgCl_2_ (Kieser et al. 2000). P-buffer containing sucrose, K_2_SO_4_, MgCl_2_, trace elements, KH_2_PO_4_, CaCl_2_, TES (Kieser et al. 2000) was used for transformation and all fusion experiments supplemented with 1 mg/mL DnaseI (Roche Diangnostics GmbH). Antibiotics apramycin (Duchefa Biochemie) and hygromycin (Duchefa Biochemie) were used for selection and were added at final concentrations of 50 μg/mL and 100 μg/mL respectively. Growth conditions for all cultures was 30°C in an orbital shaker (New Brunswick Scientific Innova®) with 100 rpm for the liquid cultures. Centrifugation (Eppendorf Centrifuge 5424) conditions were always 1000 ×g for 10 minutes (< 1 mL) or 30 minutes (>10 mL) depending on culture volume. The above mentioned culture conditions and centrifugation settings were applied throughout the study unless mentioned otherwise. All measurements for optical density of samples was done with 200 μL culture in a 96 well flat bottom plate (Sarstedt) using the Tecan spectramax platereader.

### Strain and plasmid construction

Wall deficient L-form of *Kitasatospora viridifaciens* was obtained by prolonged exposure to penicillin and lysozyme similar to a previous study (Ramijan et al. 2018). Briefly, 10^6^ spores of *Kitasatopsora viridifaciens* DSM40239 were grown in 50 mL TSBS media at 30°C and 100 rpm to obtain mycelial biomass. To this biomass 1 mg/mL lysozyme (Sigma Aldrich) and 0.6 mg/mL penicillin (Duchefa Biochemie) was added to induce S-cell formation. After 7 days, a dense culture of wall-deficient cells was obtained and subcultured to LPB media containing 6 mg/mL penicillin. This treatment was continued for 5 weeks with subculture into fresh media every week. The culture was then tested for growth on LPMA without penicillin and showed only L-form growth. A single colony was picked and inoculated in LPB without penicillin and incubated for 7 days to confirm stability of wall-deficiency and subsequently used for making a culture stock to be stored at −80°C.

The strain was further genetically modified to harbour antibiotic resistance genes and fluorescent reporter genes. Two plasmids were used for this purpose namely pGreen (containing the apramycin resistance gene *aac(3)IV* and a green fluorescent protein reporter gene) and pRed2 (containing the hygromycin resistance gene *hph* and a red fluorescent reporter gene). Both plasmids contain the Phi C31 *aatP* site and a Phi C31 integrase which allows for integration of the marker set at the *attB* site in the genome. The pGreen plasmid was obtained from a previous publication where details are provided of the construction (Zacchetti *et al*. 2016). The pRed2 plasmid was constructed by introducing the amplified mCherry gene alongwith a *gap1* promoter region at the XbaI site in the pIJ82 plasmid. Briefly, the mCherry gene was amplified together with the *gap1* promoter using primers (Sigma) mentioned in supplementary table 1 and the pRed plasmid (Zacchetti *et al*. 2016) as template. The amplified gap1-mCherry product was purified using a kit following instructions of the supplier (Illustra™ GFX™ gel band purification kit). The purified product was introduced into the vector pIJ82 at the XbaI site (New England Biolabs GmbH). This plasmid was first transformed into *E. coli* DH5alpha for amplification followed by transformation into *E. coli* ET12567 for demethylation.

The plasmids were introduced into the L-forms by polyethylene glycol (PEG1000 NBS Biologicals) induced transformation similar to protoplast transformation with some modifications (Kieser *et al*. 2000). L-form cultures were grown for 4 days. Cultures were centrifuged to remove the spent media and the pellet was resuspended in 1/4^th^ volume P-buffer. Approximately 500 ng plasmid was added to the resuspended pellet and mixed thoroughly. PEG1000 was added to this mix at a final concentration of 25 w%w and mixed gently. After a brief incubation of 5 minutes on the bench the tube was centrifuged. The supernatant was discarded and the pellet resuspended in LPB medium and incubated for 2 hours. The culture was then centrifuged again and the pellet resuspended in 100 μL LPB for plating on LPMA media containing selective antibiotics apramycin or hygromycin. After 4 days of incubation single colonies were picked and restreaked on LPMA with antibiotics for confirmation along with fluorescence microscopy. The resulting strains were named AG for Apramycin-Green and HR for Hygromycin-Red and will be referred so henceforth.

To test the antibiotic susceptibility, both strains were grown on LPMA containing with or without either 50 μg/mL apramycin or 100 μg/mL hygromycin for 4 days. Stepwise 10-fold dilution plating was done which allowed for quantifying the number of colonies (CFU/mL).

### L-form fusion

Strains AG and HR were grown individually from culture stocks in 20 mL LPB containing the relevant antibiotic. Grown cultures were then centrifuged to remove spent media containing antibiotics and washed with P-buffer twice. The pellet was finally resuspended in 2-3 mL of P-buffer containing DNase I (1 mg/mL) and the density was adjusted to 0.6 OD_600_. Both strains were then mixed in equal volumes (200 μL) in a fresh microfuge tube and mixed gently followed by incubation at room temperature for 10 minutes. Depending on the treatment, PEG1000 was added at the desired concentration (0 to 50 w%) and mixed by pipetting. For the effect of centrifugation on L-form fusion no PEG was added. After a brief incubation of 5 minutes the tubes were centrifuged and the supernatant was discarded. The pellet was resuspended in 100 μL of P-buffer with DNase I and serial dilutions were subsequently plated on LPMA with both antibiotics. Controls were also plated on the same medium such as 100 μL monocultures of each strain to test for cross resistance and 100 μL of 1:1 mix of each strain without fusion (supplementary figure 1). All plates were incubated for 3 days after which colony forming units were calculated to determine the fusion efficiency. Effiiciency was quantified as the CFU/mL on double antibiotic selection media normalized by the CFU/mL of monocultures grown on single antibiotic selection media.

### Microscopy

A Zeiss LSM 900 airyscan 2 microscope was used to image the fluorescently labeled strains under 40x magnification. For EGFP an excitation wavelength of 488 nm was used and emission captured at 535 nm whereas for mCherry an excitation wavelength of 535 nm was used and emission captured at 650 nm. Multichannel (fluorescence and brighfield), multi-stack images were captured using the Zen software (Zeiss) and further analyzed using ImageJ/Fiji. Multiple tiles were imaged for colonies to cover a large area. These tiles were then stitched and each fluorescence channel was first thresholded to determine the total pixel area. These thresholded images were then used to calculate total area (using the OR function in image calculator) and the fused area (using the AND function). The total area selection was then used to calculate individual pixel area occupied by either green or red pixels and by both.

The Lionheart FX automated microscope (BioTek) was used for timelapse imaging of double labeled L-forms after fusion. The fusant strains were precultured in LPB containing both antibiotics for 3 days. These were then centrifuged and resuspended in fresh media with antibiotics and 100 μL of this was added to individual wells in a 96 well black/clear bottom sensoplate (Thermoscientific). The plate was centrifuged for 5 minutes to enable settling of cells. The timelapse imaging was done using a 63x dry objective, set for 3 channels (brightfield, green and red) with imagining every 10 minutes for 16 hours at 30°C. The LED intensity for all channels was 10 and a camera gain of 24. The exposure time was set at the beginning of the imaging according to the reference monoculture strains AG and HR.

### Membrane fluidity assay

The membrane fluidity was quantified for cultures of different age and cultures treated with different lipopeptides using the Laurdan dye assay (Scheinpflug, Krylova, and Strahl 2017). All cultures grown in 40 mL volume were first centrifuged followed by resuspension in P-buffer and density adjusted to 0.6 to 0.8 OD_600_. The cultures were then aliquot according to the treatment for a given biological replicate (i.e. 5 aliquots of 1 mL each for 5 treatments). In case of lipopeptide treatment the lipopeptide was added to the culture at required concentration (5 μM, 10 μM or 100 μM) and all tubes were incubated for 30 minutes at 100 rpm. Centrifugation was carried out to remove excess lipopeptide and the pellet was resuspended in P-buffer. The P-buffer for this assay was always maintained at 30°C so as not to alter fluidity of the membrane. 10 mM Laurdan (6-Dodecanoyl-2-Dimethylaminonapthalene, Invitrogen) stock solution was prepared in 100% dimethylformamide (DMF, Sigma) and stored at −20°C in an amber tube to protect from light exposure. This stock solution was used to get a final concentration of 10 μM in the resuspended cultures above. The tubes were inverted to mix the dye sufficiently and then incubated at 30°C for 10 minutes and covered with foil to protect from light exposure. The cultures were then washed 3x in pre-warmed P-buffer containing 1% dimethylsulfoxide (DMSO, Sigma) to ensure removal of unbound dye molecules. The final suspension was done in pre-warmed P-buffer and 200 μL was transferred to a 96 well black/clear bottom sensoplate (Thermoscientific) for spectroscopy. Fluorescent intensities were measured by excitation at 350 nm and two emission wavelengths (435 and 490 nm). The background values were first subtracted from all sample values followed by estimation of the generalized polarization (GP) value.

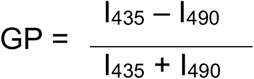

The GP value ranges from −1 to +1 with low values corresponding to high membrane fluidity.

### Lipid extraction and analysis

Cultures of the wildtype L-form were grown for different time periods (1, 3, 5 and 7 days). These were centrifuged and resuspended in P-buffer prior to membrane lipidomics. Lipids where extracted using a modified MTBE protocol of Matyash, V. et al. (ref. 10.1194/jlr.D700041-JLR200). In short, 600 μL MTBE and 150 μL methanol were added to the thawed bacteria samples. Samples where briefly vortexed, ultra-sonicated for 10 minutes and shaken at room temperature for 30 minutes. Next, 300 μL water was added and the samples where centrifuged for 5 minutes at 18213 ×g at 20 °C. After centrifugation, the upper layer was collected and transferred to a glass vial. The extraction was repeated by adding 300 μL MTBE and 100 μL methanol. Samples where briefly vortexed and shaken at room temperature for 5 minutes. Next, 100 μL water was added and the samples where centrifuged for 5 minutes at 18213 ×g at 20 °C. After centrifugation, the upper layer was collected, and the organic extracts combined. Samples where dried under a gentle stream of nitrogen. After drying samples were reconstituted in 100 μL 2-propanol. After briefly vortexing and ultra-sonication for 5 minutes, 100 μL water was added. Samples were transferred to microvial inserts for analysis

Lipidomic analysis of bacteria lipid extracts was performed using a LC-MS/MS based lipid profiling method (PMID: 31972163 DOI: 10.1016/j.bbamem.2020.183200). A Shimadzu Nexera X2 (consisting of two LC30AD pumps, a SIL30AC autosampler, a CTO20AC column oven and a CBM20A controller) (Shimadzu, ‘s Hertogenbosch, The Netherlands) was used to deliver a gradient of water:acetonitrile 80:20 (eluent A) and water:2-propanol:acetonitrile 1:90:9 (eluent B). Both eluents contained 5 mM ammonium formate and 0.05% formic acid. The applied gradient, with a column flow of 300 μL/min, was as follows: 0 min 40% B, 10 min 100% B, 12 min 100% B. A Phenomenex Kinetex C18, 2.7 μm particles, 50 × 2.1 mm (Phenomenex, Utrecht, The Netherlands) was used as column with a Phenomenex SecurityGuard Ultra C8, 2.7 μm, 5 × 2.1 mm cartridge (Phenomenex, Utrecht, The Netherlands) as guard column. The column was kept at 50 °C. The injection volume was 10 μL.

The MS was a Sciex TripleTOF 6600 (AB Sciex Netherlands B.V., Nieuwerkerk aan den Ijssel, The Netherlands) operated in positive (ESI+) and negative (ESI−) ESI mode, with the following conditions: ion source gas 1 45 psi, ion source gas 2 50 psi, curtain gas 35 psi, temperature 350°C, acquisition range *m/*z 100-1800, ion spray Voltage 5500 V (ESI+) and −4500 V (ESI−), declustering potential 80 V (ESI+) and −80 V (ESI−). An information dependent acquisition (IDA) method was used to identify lipids, with the following conditions for MS analysis: collision energy ±10, acquisition time 250 ms and for MS/MS analysis: collision energy ±45, collision energy spread 25, ion release delay 30, ion release width 14, acquisition time 40 ms. The IDA switching criteria were set as follows: for ions greater than *m/z* 300, which exceed 200 cps, exclude former target for 2 s, exclude isotopes within 1.5 Da, max. candidate ions 20. Before data analysis, raw MS data files were converted with the Reifycs Abf Converter (v1.1) to the Abf file format. MS-DIAL (v4.20), with the FiehnO (VS68) database was used to align the data and identify the different lipids (Tsugawa et al. 2015; 2019; 2020). Further processing of the data was done with R version 4.0.2 (R Core Team 2014).

The relative abundance of specific lipid class *vs* total relative abundance was used to roughly compare the ratio of each lipid class. The lipids have been sorted into saturated and unsaturated lipids classes. Also, the lipids have been sorted based on head groups (DG, TG, PE, PI) and the ratio of each class have been calculated

### Lipopeptide preparation and treatment

Peptide K_4_ and E_4_ were synthesized on a CEM Liberty Blue microwave-assisted peptide synthesizer using Fmoc chemistry. 20% piperidine in DMF was used as the deprotection agent. During coupling, DIC was applied as the activator and Oxyma as the base. All peptides were synthesized on a Tentagel S RAM resin (0.22 mmol/g). The resin was swelling for at least 15min before synthesis started. For the coupling, 5 equivalents of amino acids (2.5 mL in DMF), DIC (1 mL in DMF) and Oxyma (0.5 mL in DMF) were added to the resin in the reaction vessel and were heated to 90 °C for 4 minutes to facilitate the reaction. For deprotection, 20% of piperidine (4 mL in DMF) was used and heated to 90°C for 1 minute. Between deprotection and peptide coupling, the resin has been washed three times using DMF. After peptide synthesis, a polyethyleneglycol (PEG)_4_ linker and cholesterol were coupled manually to the peptide on-resin. 0.1 mmol of each peptide was reacted with 0.2 mmol N_3_-PEG_4_-COOH by adding 0.4 mmol HCTU and 0.6 mmol DIPEA in 3 mL DMF. The reaction was performed at room temperature for 5 hours. After thorough washing, 3 mL of 0.5 mmol trimethylphosphine in a 1,4-dioxane:H_2_O (6:1) mixture was added to the resin to reduce the azide group to an amine (overnight reaction). After reduction, the peptide was reacted with cholesteryl hemisuccinate (0.3 mmol) in DMF by adding 0.4 mmol HCTU and 0.6 mmol DIPEA. The reaction was performed at room temperature for 3 hours. Lipopeptides were cleaved from the resin using 3 mL of a TFA:triisopropylsilane (97.5:2.5%) mixture and shaking for 50 min. After cleavage, the crude lipopeptides were precipitated by pouring into 45 mL of −20 °C diethyl ether:n-hexane (1:1) and isolated by centrifugation. The pellet of the lipopeptides was redissolved by adding 20 mL H_2_O containing 10% acetonitrile and freeze-dried to yield a white powder. Lipopeptides were purified with reversed-phase HPLC on a Shimazu system with two LC-8A pumps and an SPD-20A UV-Vis detector, equipped with a Vydac C4 column (22 mm diameter, 250 mm length, 10 μm particle size). CPK_4_ was purified using a linear gradient from 20 to 65 % acetonitrile in water (with 0.1% TFA) with a 12 mL/min flow rate over 36 mins. CPE_4_ was purified using a linear gradient from 20 to 75 % acetonitrile in water (with 0.1% TFA) with a 12 mL/min flow rate over 36 mins. After HPLC purification, all peptides were lyophilized and yielded white powders. For the fluo-K_4_ and fluo-E_4_ synthesis, two additional glycine residues were coupled to the N-terminus of the peptides on resin, before the dye was manually coupled by adding 3 mL DMF containing 0.2 mmol 5(6)-carboxyfluorescein, 0.4 mmol HCTU and 0.6 mmol DIPEA. The reaction was left at room temperature overnight. The fluo-K_4_ and fluo-E_4_ were cleaved from the resin using 3 mL of a TFA:triisopropylsilane:H_2_O (97.5:2.5%) mixture and shaking for 1.5 hours. After cleavage, the crude lipopeptides were precipitated by pouring into 45 mL of −20 °C diethyl ether and isolated by centrifugation. The pellet of the lipopeptides was redissolved by adding 20 mL H_2_O containing 10% acetonitrile and freeze-dried to yield a white powder. Fluo-K_4_ and fluo-E_4_ were purified using the same HPLC described above equipped with a Kinetix Evo C18 column (21.2 mm diameter, 150 mm length, 5 μm particle size). For the fluo-K4, a linear gradient from 20 to 45% acetonitrile in water (with 0.1% TFA) with a 12 mL/min flow rate over 28 mins was used. For fluo-E_4_, linear gradient from 20 to 55% was used. After HPLC purification, all peptides were lyophilized and yielded orange powders. The purity of all peptides were determined by LC-MS (supplementary table 2). The structure of all peptides used in this study can be found in supplementary figure 3. Treatment of cultures with different peptides was done by adding externally to cells suspended in P-buffer and incubating for 30 minutes at 30°C 100 rpm. Excess peptide was washed by centrifugation.

### L-form membrane labelling

3×10^8^ wild type L-forms were suspended in 1 mL of P-buffer. 10 μL of CPK_4_ or CPE_4_ (10 mM in DMSO) was added to the L-form suspension to a final concentration of 100 μM. After 30 min incubation at 30 °C with shaking at 100 rpm, the L-forms were washed two times by centrifugation using P-buffer. The L-forms were then suspended in 900 μL P-buffer and 100 μL of fluo-K_4_ or fluo-E_4_ (200 μM in P-buffer) was added to a final concentration of 20 μM. After 5min incubation, the L-forms were washed three times using P-buffer to get rid of the free fluorescent lipopeptides. For control experiments, fluo-K_4_ or fluo-E_4_ were added to non-lipopeptide modified L-form and incubated for 5 min. L-form imaging was performed on a Leica SP8 confocal microscopy. Excitation: 488 nm, emission: 500-550 nm.

### Peptide induced L-form fusion

Strains AG and HR were grown individually from culture stocks in 20 mL LPB containing the relevant antibiotic. Grown cultures were then centrifuged to remove spent media containing antibiotics and washed with P-buffer twice. The pellet was finally resuspended in 2-3 mL of P-buffer containing DNase I (1 mg mL^−1^) and the density was adjusted to 0.6 OD_600_. Peptides were added at required concentrations to 1 ml cultutres of individual strains AG and HR. Cultures were then incubated for 30 minutes at 30 °C with shaking at 100 rpm. Excess and unbound peptide was removed via centrifugation and resuspension of pellet in 1 ml P buffer containing DNase I. Both strains were then mixed in equal volumes (200 μL) in a fresh microfuge tube and mixed gently followed by incubation at room temperature for 10 minutes. Depending on the treatment cultures were centrifuged followed by treatment with PEG1000 or simply centrifuged. The pellet was resuspended in 100 μL of P-buffer with DNase I and serial dilutions were subsequently plated on LPMA with both antibiotics. Controls were also plated on the same medium such as 100 μL monocultures of each strain to test for cross resistance and 100 μL of 1:1 mix of each strain without fusion (supplementary figure 1). All plates were incubated for 3 days after which colony forming units were calculated to determine the fusion efficiency. Effiiciency was quantified as the CFU/mL on double antibiotic selection media normalized by the CFU/mL of monocultures grown on single antibiotic selection media.

### Statistical analysis and graphs

Statistical analysis of all datasets was done in R version 3.6.1 (R Core Team 2014) using built-in packages. The specific tests performed are mentioned in the results and figure legends. All graphs were produced using the package ggplot2 (Wickham, H. 2009).

## Acknowledgements

We thank members of the Claessen lab and Kros lab for fruitful discussions and suggestions. S.S. acknowledges the NWA startimpulse (Origins Centre) for funding.

## Author contribution

S.S., D.C, and A.K. designed the project. S.S. performed all experiments. S.S. and M.S. performed peptide fusion experiments. M.S. prepared all lipopeptides and did microscopy for lipopeptide docking experiments. B.C. prepared the cell-wall-deficient line of *K. viridifaciens* used in the study. R.D. and M.G. performed the membrane lipid analysis. S.S., D.R., D.C. and A.K. acquired funding. S.S. wrote the first draft followed by revisions from all authors. All authors approved the final manuscript.

