## Supplementary figures and table for "Generating heterokaryotic cells via bacterial cell-cell fusion"


| Supplementary | Page no. |
|---|---|
| Figures | 2-6 |
| Tables | 7 |

#### Supplementary figure 1 (to Figure 3)

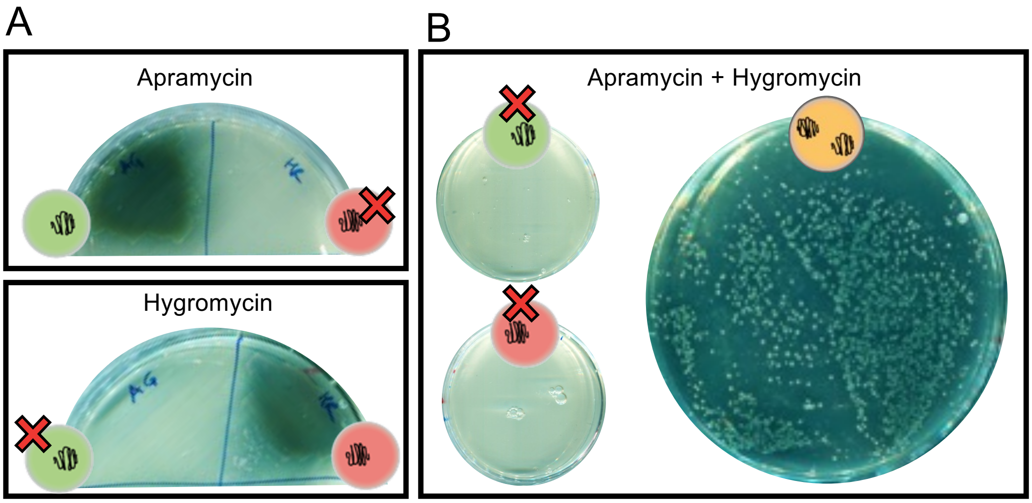


**Confirmation of phenotypes.** (**A**) Strains AG and HR plated on single antibiotic (either apramycin or hygromycin) to confirm resistance prior to fusion. AG shows growth on apramycin (top) and HR on hygromycin (bottom). (**B**) After fusion, monocultures of AG and HR plated on media with both antibiotics show no growth. Cells that have undergone fusion grow in the presence of both apramycin and hygromycin.

#### Supplementary figure 2 (for figure 4)

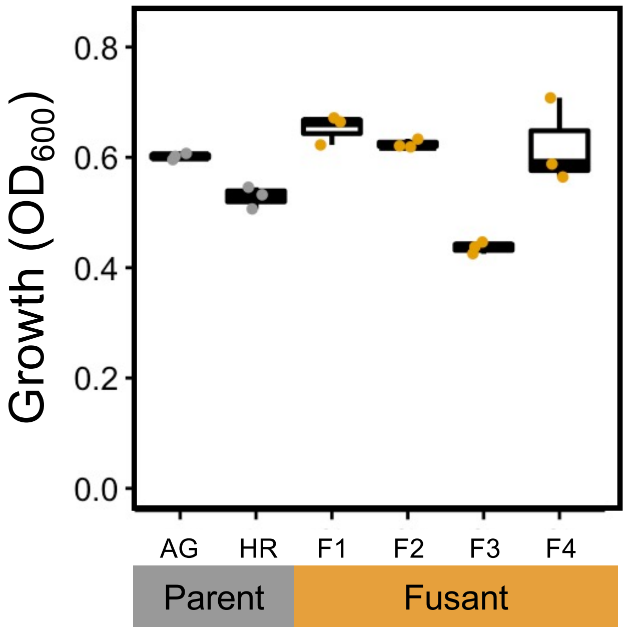


**Fused cells show growth comparable to parent strains.** All cultures; parent (grey) and fusant (yellow) were grown in liquid media under selection (apramycin for AG, hygromycin for HR and both for the fusant) for 4 days at 30ºC to confirm viability after fusion. Optical density measured at 600 nm indicates growth of all populations.

### Supplementary figure 3 (for figure 6)

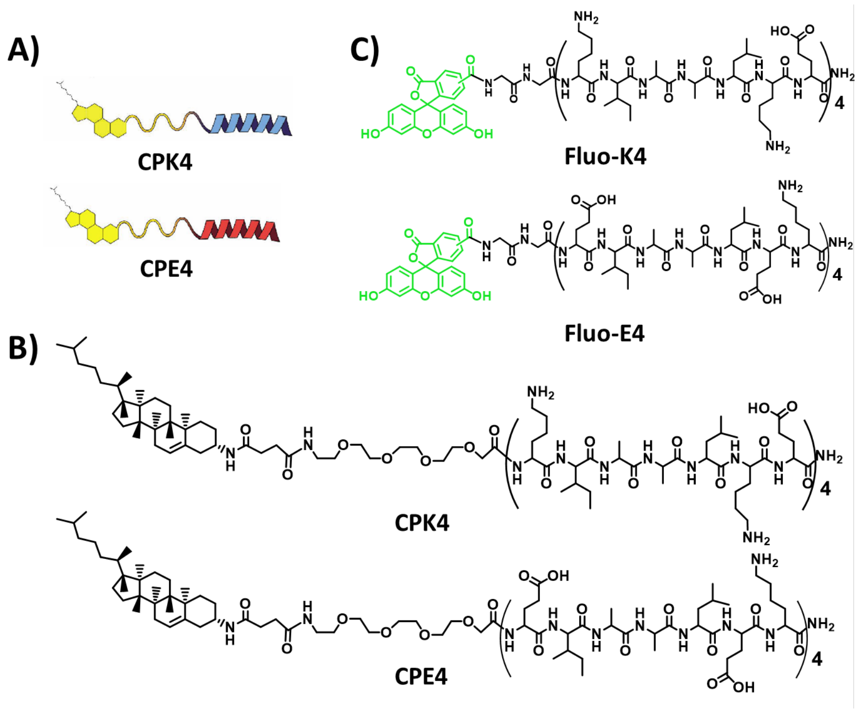


**Schematic representation** (A) and chemical structure (B) of $CPK_4$ and $CPE_4$. (C) Chemocal structure of fluo-$K_4$ and fluo-$E_4$.

Supplementary figure 4 (for figure 6)

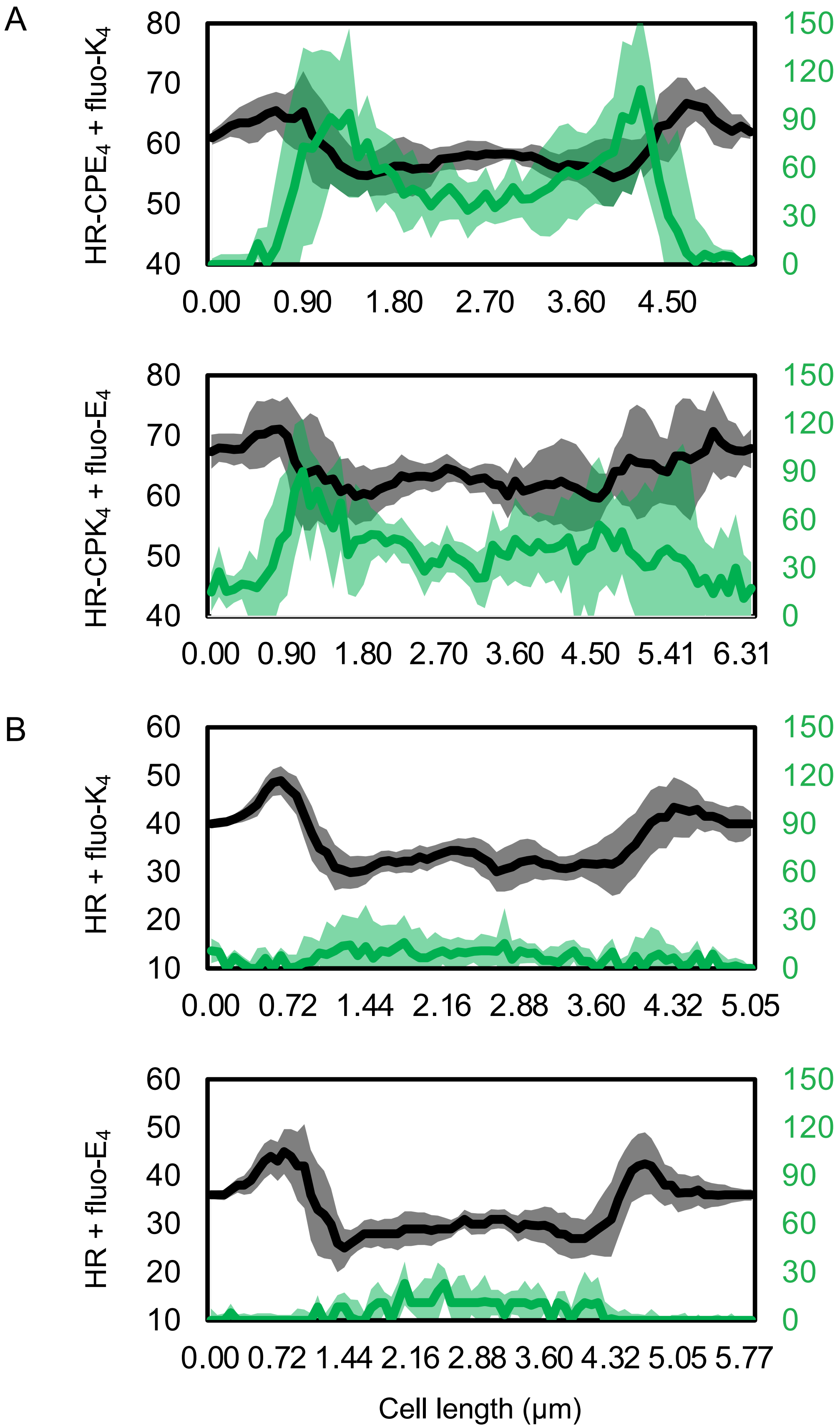

**Coiled coil peptides localize on the membrane.** (**A**) Image analysis of the membrane labelled L-forms (HR) was done to obtain intensity profiles across the cell (n=10). Shown is the median ± SD values for the cell in brightfield (black, left y axis) and the eGFP (green, right y axis) for the different treatments (same as in figure 3). The profiles indicate peaks of fluorescence (green) coinciding with the boundary of cells also seen as peaks (black). The fluorescence intensities decrease in the central region of the cell. (**B**) The controls where no coiled coil lipopeptide ($CPE_4$ or $CPK_4$) was added to the cells show no peaks of fluorescence (green) across the cell profile (black).

Supplementary figure 5 (to Figure 7)

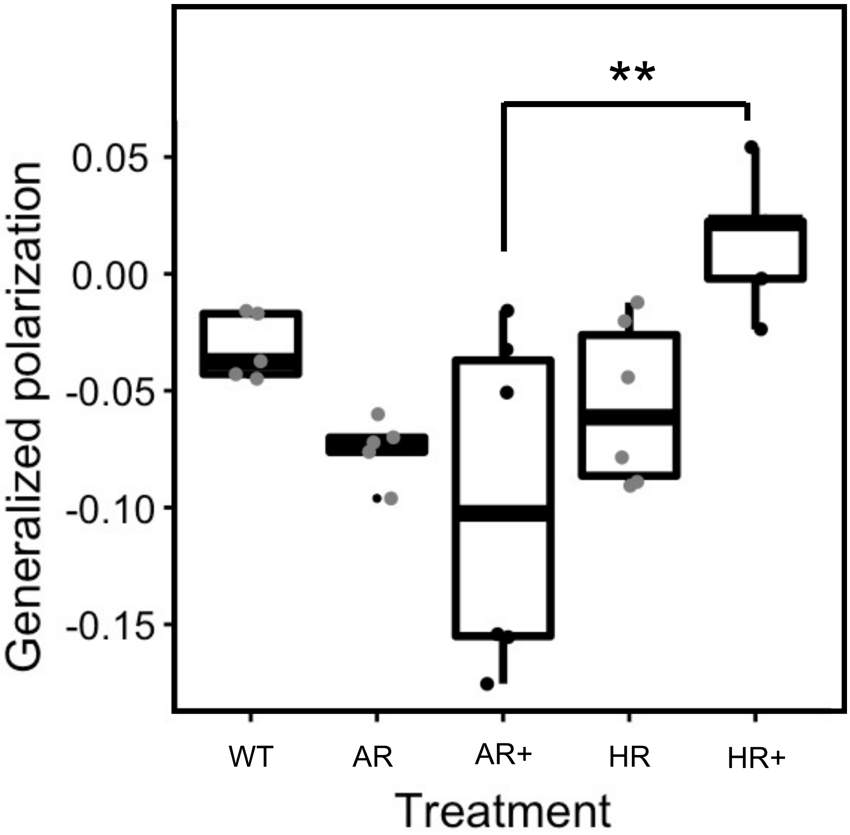


**Antibiotic alters membrane fluidity.** Strains (WT, AR and HR) were grown in LPB media in the absence or presence (+) of the selective antibiotic (apramycin for AR and hygromycin for HR) for 2 days and tested for membrane fluidity using the Laurdan assay. WT, AR and HR have similar basal levels of fluidity around -0.05. Apramycin treatment increases the fluidity of the AR strain whereas hygromycin treatment decreases fluidity of the HR strain. These differences in GP value between the AR+ and HR+ conditions (One-way ANOVA, F=5.85, p=0.002 followed by Tukey's pairwise comparison) are similar to that observed in figure 6 in the control treatment of AR and HR strains.

### Supplementary figure 6 (for figure 7)

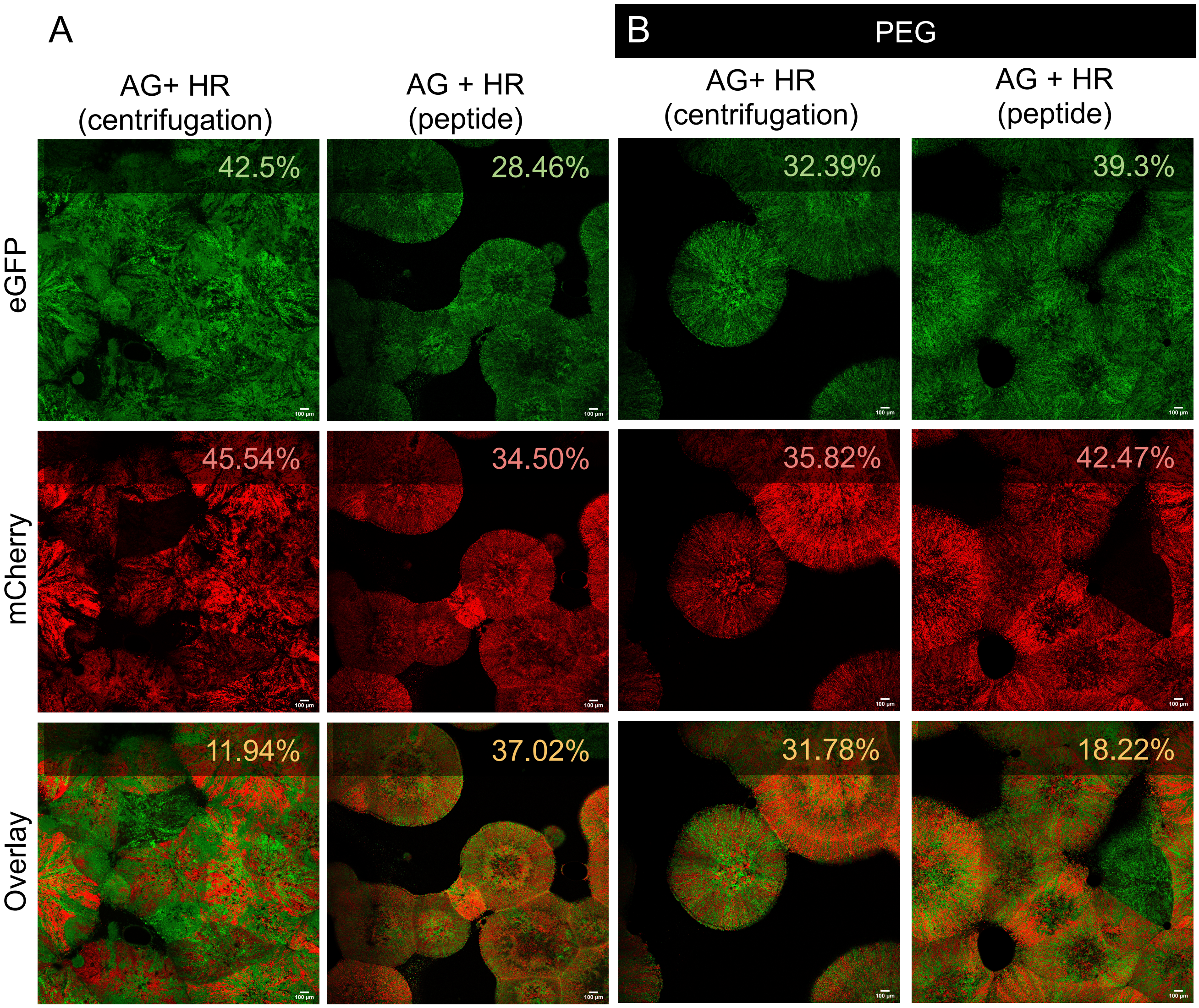


**Microscopy of fused colony.** (**A**) Strains AG and HR were fused using centrifugation and peptides followed by selection of colonies on double antibiotic medium. Whole colonies were imaged for measuring expression of fluorescence reporters EGFP (top row) and mCherry (middle row). The percent of fluorescence is indicated in the top right corner of each image and was calculated using ImageJ/Fiji. (**B**) Fusion of strains using centrifugation or peptides in combination with PEG resulted in more colonies on double selection medium. This colonies were imaged and fluorescence percent was calculated. Scale bar = 100 µm.

**Supplementary table 1.** Primer sequences used to amplify gap1-mCherry for the pRed2 plasmid construction.

| **Primer** | **Sequence 5'-3'** |
|---|---|
| Gap-FW-XbaI | GATTACTCTAGACCGAGGGCTTCGAGAC |
| mCherry-RV-XbaI | TAAGCATCTAGACTAGCCGCCACACTTGTAC |

**Supplementary table 2.** Calculated mass and found mass via LC-MS of $CPK_4$ and $CPE_4$.

| **Peptide** | **Mass (calcd.) / Da** | **Mass (found) / Da** |
|---|---|---|
| **$CPK_4$** | $[M + 2H^+]^{2+}$ 1867.9 | 1866.4 |
| | $[M + 3H^+]^{3+}$ 1245.2 | 1244.3 |
| **$CPE_4$** | $[M + 2H^+]^{2+}$ 1869.7 | 1868.8 |
| | $[M + 3H^+]^{3+}$ 1246.5 | 1245.4 |
| **Fluo-$K_4$** | $[M + 2H^+]^{2+}$ 1876.5 | 1874.0 |
| | $[M + 3H^+]^{3+}$ 1251.3 | 1249.3 |
| **Fluo-$E_4$** | $[M + 2H^+]^{2+}$ 1877.4 | 1875.9 |
| | $[M + 3H^+]^{3+}$ 1251.6 | 1250.6 |
